# Sequential Development of Task Representation from Hippocampus to Prefrontal Cortex Supports Goal-Directed Spatial Navigation

**DOI:** 10.1101/2025.10.05.680599

**Authors:** Heung-Yeol Lim, Sewon Park, Inah Lee

## Abstract

Successful goal-directed navigation requires the coordination between the hippocampus and medial prefrontal cortex. However, it is not fully confirmed that the medial prefrontal cortex learns its spatial code from the hippocampus. To test this, we examined spatial representations of the intermediate hippocampus and medial prefrontal cortex while rats learned a spatial navigation task. Rats performed a goal-directed spatial navigation task in 2D VR to find an unmarked goal zone, and we discovered robust directional tuning of single neurons in both regions. Neural manifold analysis further confirmed population-level directional tuning in both regions, with manifolds having ring-like geometry. We found that this ring-like structure evolved after learning, in a way that the hippocampal-prefrontal manifolds converged to a shared geometry. Furthermore, this evolution of ring-like structure was preceded by the hippocampus at the trial level. It was further verified that the evolution of the ring-like structure is linked to phase locking to the hippocampal theta rhythm, particularly in the prefrontal manifolds. Our findings provide compelling evidence that spatial representations of the hippocampal-prefrontal network become aligned after learning, and also highlight the information flow from the hippocampus to the medial prefrontal cortex during this geometry synchronization.

## Introduction

Successful navigation relies on an internal representation of the external world, the cognitive map, encoded by hippocampal place cells^1–3^. Converging evidence indicates that the hippocampus works in concert with the medial prefrontal cortex (mPFC), a region implicated in planning, decision making, and schema formation^4–8^, to facilitate navigation^9–13^. In line with this account, lesions or inactivations of the hippocampal-prefrontal networks produce marked deficits in various spatial navigation tasks, especially when their connections were disrupted by contralateral silencing^14–18^. Moreover, the mPFC contains spatially modulated neurons, similar to hippocampal place cells^19–22^, and its spatial coding is perturbed after disrupting the hippocampus^23,24^.

Building on this foundation, studies have highlighted physiological coupling between the hippocampus and mPFC during spatial navigation, particularly via theta oscillations originating in the hippocampus^25–28^. For example, in a spatial working memory task, hippocampal-prefrontal theta coherence and phase-locked spiking were enhanced in correct trials requiring higher memory demands^25^. This theta-based coupling is also tied to spatial coding in the mPFC, as shown by enhanced spatial decoding from the mPFC population after incorporating hippocampal theta rhythm^21^. Together, these findings emphasize the importance of hippocampal-prefrontal coupling for successful navigation, yet most work has examined well-trained animals in post-learning states. As a result, how hippocampal and prefrontal spatial representations dynamically emerge and interact during learning remains poorly understood, raising the question of whether the mPFC acquires its spatial code from the hippocampus.

In addition, it should be noted that most prior work on spatial navigation focused on the dorsal hippocampus, although its connections to mPFC rely on indirect pathways via other thalamic and cortical regions^29–31^. The ventral hippocampus (including the intermediate part) projects monosynaptically to the mPFC. However, the most ventral part of the hippocampus exhibits weak positional firing^29,30,32^ and its major roles have been implicated in anxiety-like behavior and novelty processing^33–35^. In this regard, the intermediate hippocampus (iHP) occupies a unique position in communicating spatial codes to the mPFC because it has direct connectivity to the mPFC while retaining spatial precision^36,37^. Previous work from our laboratory has further demonstrated that, after the reward values of two locations were reversed, iHP place cells immediately remapped to represent the high-value location, while dorsal hippocampus place cells maintained their preferred location^36,38^. These results highlight the iHP’s unique role in integrating spatial and value-based motivational information^36,39^, and this value-associated spatial map can be a plausible source for goal-directed actions by the mPFC.

In the current study, with simultaneous iHP and mPFC recordings, we tested whether the mPFC learns its spatial code from iHP to support successful spatial navigation. In our experiment, rats learned a goal-directed spatial navigation task in a two-dimensional (2D) virtual reality (VR) environment. We directly compared directional tuning, a prominent spatial signal in 2D VR space, before and after learning the task^40,41^. We observed post-learning refinement of directional tuning at the single-neuron and population level (i.e., manifold geometry) in both regions. We also demonstrated a sequential development of directional representations from the iHP to mPFC by analyzing trial-by-trial dynamics. Lastly, we examined the relationship between theta synchronization and directional tuning and discovered that iHP-to-mPFC theta transmission was associated with directional refinement in the mPFC. Taken together, our findings provide compelling evidence that iHP provides a spatial map to the mPFC over the course of learning the spatial task to guide successful navigation.

## Results

### Rats successfully navigate a 2D space to find a goal location in a VR environment

We employed a 2D VR navigation system for rodents, as previously described^39^ (**Fig. 1A**). In this experimental setup, rats were body-restrained while they navigated a circular arena by rotating a spherical treadmill. The rats were trained to perform a goal-directed spatial navigation task using distal visual landmarks in the VR environment (**Fig. 1B, C**). Two unmarked goal zones were positioned on the west and east sides of the arena (**Fig. 1B**). At the onset of each trial, rats were placed at the center of the arena, facing north or south, with the starting orientation predetermined pseudorandomly. Initially, rats were trained to navigate to the western goal zone to obtain a water reward (West condition). Trials in which rats entered the opposite zone (e.g., East) were immediately terminated without reward and recorded as incorrect. Upon the rats reaching ≥ 75% accuracy (chance = 50%) for two consecutive days, the rewarded location was switched to the eastern goal zone (East condition), which the rats learned to navigate to by trial and error.

**Fig. 1.**
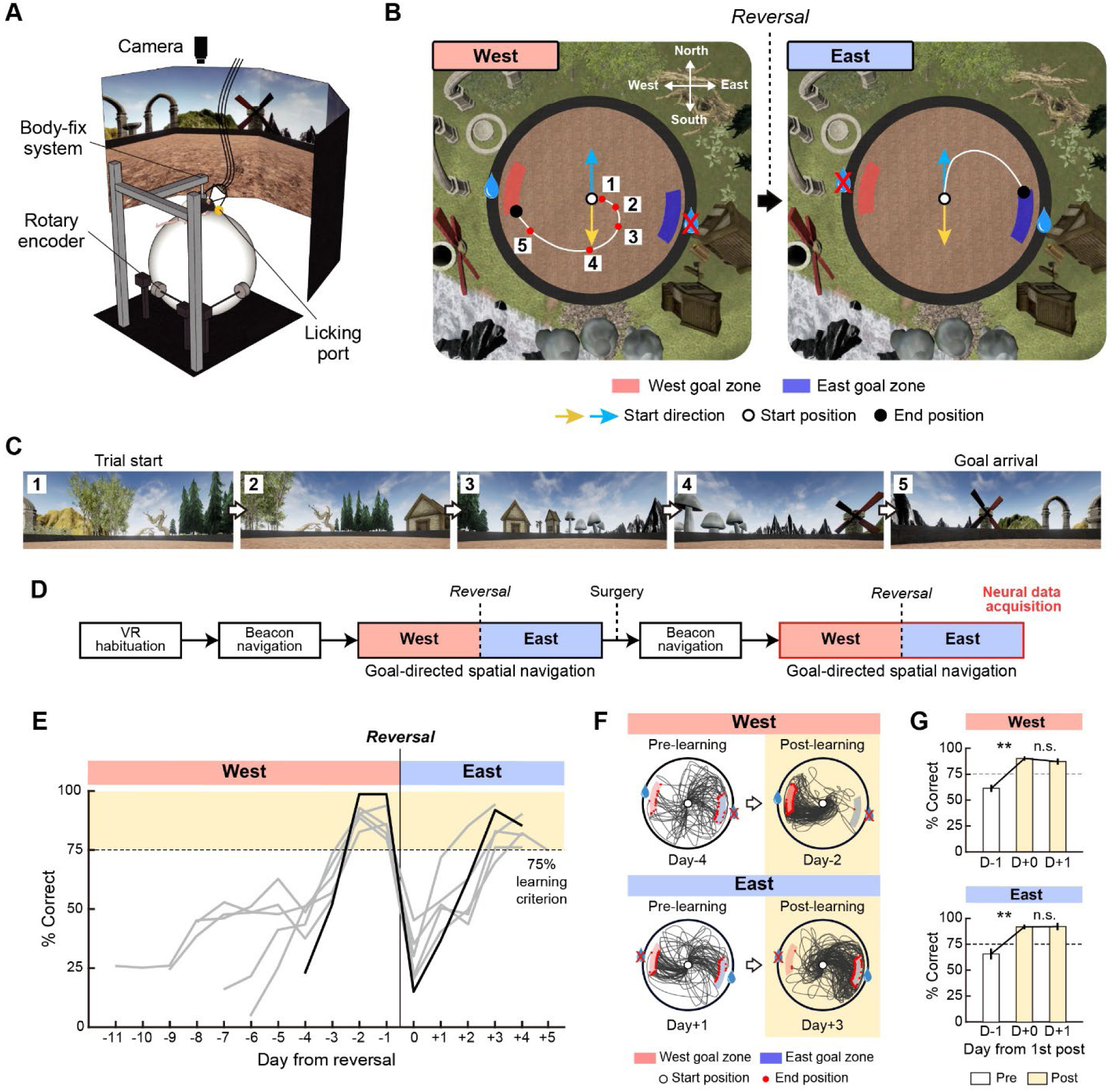
Goal-directed spatial navigation task in a 2D VR environment. (**A**) 2D VR apparatus. (**B**) Bird’s-eye view of the VR space and example trajectories during West and East conditions. Rats initially learned to navigate to the West goal zone to get a water reward. After reaching a learning criterion, the goal zone was reversed to the East. Start direction (north or south) was pseudorandomly assigned across trials. The goal zones were unmarked. (**C**) Visual scenes surrounding the VR environment. Numbers in the top-left corner indicate the rat’s position marked in **B**. (**D**) Training and recording schedule for the spatial navigation task. (**E**) Across-day behavioral performance of all rats (n = 6). Sessions with less than 75% accuracy were defined as the pre-learning stage, while those more than 75% were defined as the post-learning stage (highlighted in yellow). The black line indicates the performance of an example rat whose trajectories are shown in **F**. (**F**) Example trajectories of the rat during pre- and post-learning in the West and East conditions. (**G**) Average accuracy of all rats, aligned to the first day exceeding 75% accuracy (dotted line). Data represent the mean ± SEM. **p < 0.01; n.s., not significant.

All rats (n = 6) successfully learned the task in both the West and East conditions, although the number of days required to meet the learning criterion varied among rats (**Fig. 1E**). To compare behavior and neural activity before and after task learning, we designated sessions with fewer than 75% correct trials as ‘pre-learning’ and those with equal to or more than 75% as ‘post-learning’ (**Fig. 1E, F**). There were significant performance differences between pre- and post-learning sessions (West: F_(2,10)_ = 24.42, p = 0.004; East: F_(2,10)_ = 26.53, p = 0.004; one-way repeated measures ANOVA) (**Fig. 1G**). On the first day that rats surpassed the 75% accuracy threshold (i.e., the first post-learning day), performance increased sharply relative to the preceding day (West, t_(5)_ = 6.19, p = 0.003; East, t_(5)_ = 5.87, p = 0.004; paired t-test with Bonferroni correction [n = 2]), indicating a distinct transition in learning (**Fig. 1G**). Performance remained consistently high on the subsequent day (i.e., the second post-learning day) (West, t_(5)_ = 1.26, p = 0.52; East, t_(5)_ = 0.095, p = 1; paired t-test with Bonferroni correction [n = 2]) (**Fig. 1G**).

### Robust directional tuning of iHP and mPFC neurons in 2D VR space, with higher precision in iHP

We recorded single-unit spiking simultaneously from the iHP and the mPFC during pre- and post-learning stages of the spatial navigation task (**Fig. 2A, C**). To investigate spatial coding at the single-neuron level, we analyzed putative excitatory neurons in the iHP (n = 1228) and the mPFC (n = 410), focusing on directional tuning, which is known to be prominent in visually enriched 2D VR environments^40,41^. VR direction was defined as the allocentric orientation of the virtual agent in the VR space (**Fig. 2B**), and firing rate variations were quantified as a function of the VR direction.

**Fig. 2.**
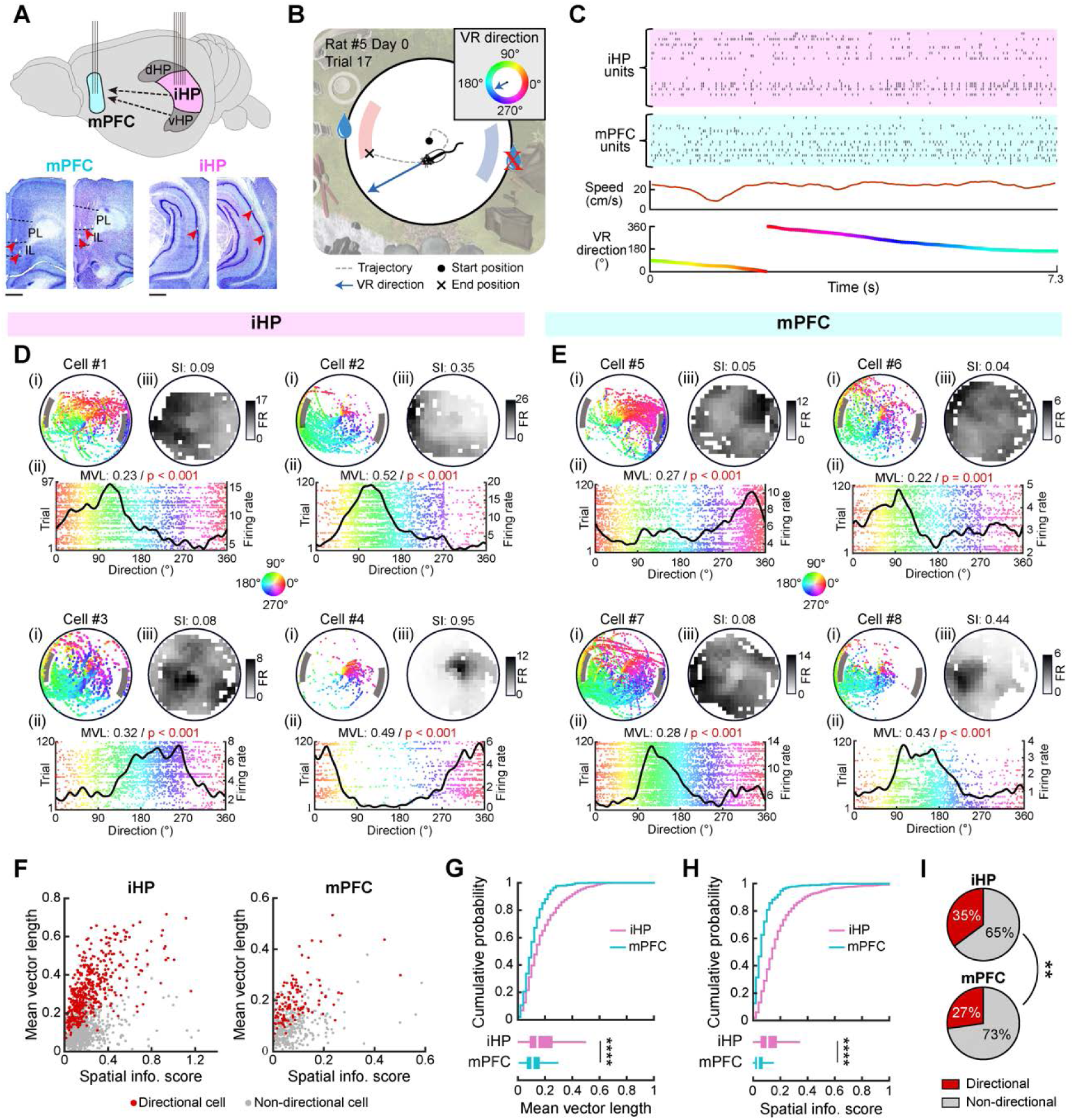
Directional tuning in the iHP and mPFC during 2D VR navigation. (**A**) Schematic of simultaneous recordings in the iHP and the mPFC (top). The dotted arrows highlight direct projections from the intermediate/ventral hippocampus (i/vHP) to the medial prefrontal cortex (mPFC). Histological verification of the recording sites (bottom). Red arrowheads indicate tetrode tip locations. PL, prelimbic; IL, infralimbic. Scale bar indicates 0.5 mm. (**B**) Definition of VR direction during 2D VR navigation. (**C**) Simultaneously recorded spike trains from the iHP and the mPFC, with corresponding behavioral measures (speed and VR direction), during the example trial shown in **B**. (**D**, **E**) Representative examples of single neurons exhibiting directionally selective firing in the iHP and the mPFC, shown as (i) directional spike maps, (ii) directional tuning curves with raster plots, and (iii) spatial firing rate maps. Spike colors indicate VR direction. MVL, mean vector length; SI, spatial information score. P-values were obtained by comparing MVL to surrogate data. (**F**) Scatter plots of mean vector length versus spatial information score for directional and non-directional cells in the iHP and the mPFC. (**G**) Comparison of mean vector length for directional tuning between the iHP and the mPFC, shown as cumulative distributions (top) and box plots (bottom). (**H**) Same as in **G**, but for spatial information score. (**I**) Proportion of directional cells in the iHP and mPFC. **p < 0.01, ****p < 0.0001.

Analysis of spike distributions relative to VR direction revealed robust directional tuning in both iHP (**Fig. 2D**) and mPFC (**Fig. 2E**). For example, in figure 2D, cell #1 fired selectively near 90°, as evidenced by its spike map (**Fig. 2D-i**) and directional tuning curve (**Fig. 2D-ii**). Neurons were classified as directionally tuned based on the significance of mean vector length compared to the surrogate distribution (α = 0.05). To account for potential covariance between direction and position during goal-directed navigation (see trajectory examples in **Fig. 1F**), we assessed positional selectivity using spatial rate maps and spatial information scores (**Fig. 2D-iii, 2E-iii**). While some neurons exhibited both directional and positional selectivity (e.g., iHP cell #4; mPFC cell #8), many directionally tuned neurons fired broadly across the arena with low spatial information (e.g., iHP cells #1–3; mPFC cells #4–7). Across the population, a substantial proportion of neurons displayed high MVL despite low spatial information, suggesting that directional tuning was not solely attributable to positional encoding (**Fig. 2F**).

Comparative analysis revealed that the iHP neurons exhibited significantly higher MVL than mPFC neurons (z = 8.16, p < 0.0001; Mann-Whitney U test) (**Fig. 2G**). In addition, the proportion of directionally tuned cells was significantly higher in the iHP than in the mPFC (χ^2^ = 8.57, p = 0.003; chi-square test) (**Fig. 2I**). Spatial information scores were also significantly higher in the iHP compared to the mPFC (z = 17.14, p < 0.0001; Mann-Whitney U test) (**Fig. 2H**). Taken together, these findings confirm robust directional tuning in both the iHP and the mPFC in the 2D VR environment, consistent with prior reports in the dorsal hippocampus^40,41^, and indicate that such tuning is not primarily driven by positional coding. Moreover, the iHP displayed greater specificity in directional tuning compared to the mPFC, consistent with prior studies reporting weaker spatial representations in the prefrontal cortex^19,21,42^.

### Post-learning refinement of directional tuning in both the iHP and the mPFC, with more predominant effects in the mPFC

We investigated how directional tuning evolves with learning. Example neurons demonstrated sharper directional tuning in post-learning sessions, as indicated by increased MVL compared to pre-learning sessions (**Fig. 3A, B**). At the population level, MVL increased significantly from pre-learning to post-learning in both regions (F_(1,540)_ = 18.16, p < 0.0001, two-way ANOVA; iHP-Pre vs. iHP-Post, t_(430)_ = 3.14, p = 0.0036; mPFC-Pre vs. mPFC-Post, t_(110)_ = 4.81, p < 0.0001; t-test with Bonferroni correction [n = 2]), with the iHP maintaining significantly higher MVL than the mPFC in both stages (F_(1,540)_ = 97.81, p < 0.0001; two-way ANOVA) (**Fig. 3C**). The proportion of directionally tuned neurons was greater in the iHP during pre-learning (χ^2^ = 18.76, p < 0.0001), but this regional difference diminished in post-learning sessions due to a significant increase in the proportion of directionally tuned neurons in the mPFC (χ^2^ = 0.51, p = 0.47; chi-square test) (**Fig. 3D**). These results suggest that directional tuning strengthens with learning in both regions, with the mPFC exhibiting more pronounced learning-related changes.

**Fig. 3.**
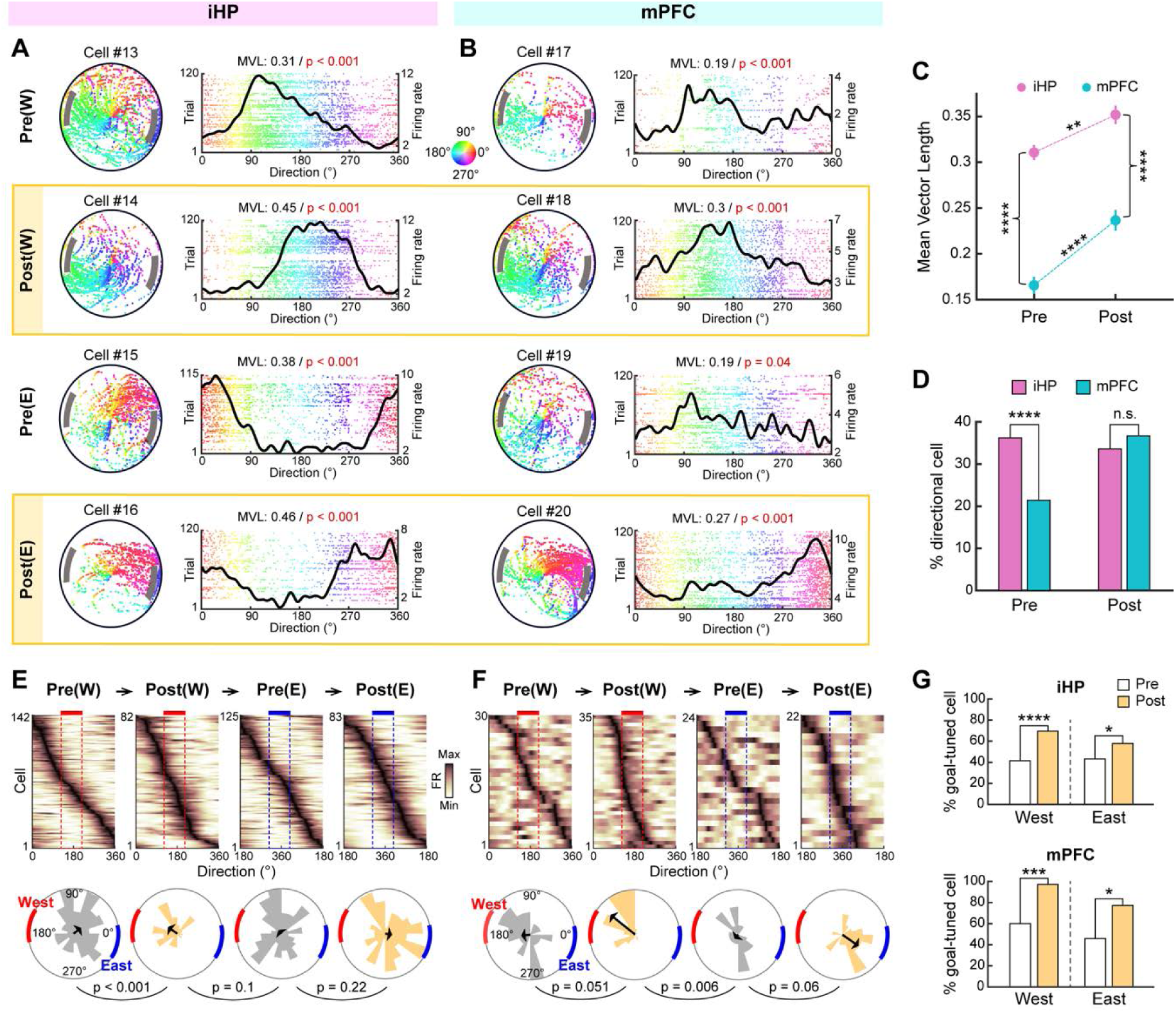
Post-learning refinement of directional tuning in the iHP and the mPFC. (**A**, **B**) Representative examples of directional cells in the iHP (**A**) and the mPFC (**B**) in the pre- and post-learning stages of West (W) and East (E) conditions. Spike colors indicate VR direction. MVL, mean vector length. (**C**) Comparison of mean vector length for directional cells in the iHP and mPFC during pre- and post-learning stages. Data represent the mean ± SEM. (**D**) Proportions of directional cells in the iHP and the mPFC during pre- and post-learning stages. (**E**, **F**) Population of directional tuning curves aligned to their preferred directions (top) in the iHP (**E**) and the mPFC (**F**) across learning stages. Dotted squares indicate a 90° range encompassing the center of the goal-zone direction (170° in West [red] and 350° in East [blue] conditions). Rose plots (bottom) illustrate the distribution of preferred directions with arrows indicating mean vectors. P-values were computed using Watson’s U2 test with Bonferroni correction (n = 3). (**G**) Proportions of goal-tuned cells preferring the goal-zone direction in the iHP and mPFC during pre- and post-learning stages, separately for West and East conditions. *p < 0.05, **p < 0.01, ***p < 0.001, ****p < 0.0001; n.s., not significant.

To assess whether directional tuning reflects the current reward location, we analyzed directional tuning curves aligned to each neuron’s peak direction for West and East conditions across pre- and post-learning stages (**Fig. 3E, F**). In both the iHP (**Fig. 3E**) and the mPFC (**Fig. 3F**), directional tuning spanned the full 360°, but post-learning sessions showed an overrepresentation of directions near the reward location (170° for West; 350° for East). Circular statistics supported goal-biased representations during post-learning in the iHP (Pre(W) vs. Post(W), p < 0.001; Post(W) vs. Pre(E), p = 0.1; Pre(E) vs. Post(E), p = 0.22) and particularly in the mPFC (Pre(W) vs. Post(W), p = 0.051; Post(W) vs. Pre(E), p = 0.006; Pre(E) vs. Post(E), p = 0.06; Watson’s U2 test with Bonferroni correction [n = 3]). The proportion of goal-tuned cells, defined as those with preferred directions within the range facing the current reward location, significantly increased from pre-learning to post-learning in both regions (iHP-West, χ^2^ = 16.28, p < 0.0001; iHP-East, χ^2^ = 4.27, p = 0.039; mPFC-West, χ^2^ = 14.33, p = 0.0002; mPFC-East, χ^2^ = 4.76, p = 0.029; chi-square test) (**Fig. 3G**). These findings indicate that learning not only sharpens directional tuning but reorients it toward goal-relevant directions, facilitating effective goal-directed navigation.

### Evolution of directional neural manifolds after learning, with a stronger ring-like structure in the iHP

To explore population-level representations, we employed neural manifold analysis to examine the geometry of population activity in the iHP and the mPFC (**Fig. 4A, B**). Directional tuning curves (5° bins; 72 bins covering 360°) were constructed from simultaneously recorded neurons in a session from the iHP or the mPFC and projected into three-dimensional space using nonlinear dimensionality reduction (Isomap). Each manifold consists of 72 points, capturing the neural state of the population at each direction bin. In both the iHP and the mPFC, the resulting manifolds typically exhibited a ring-like structure, reflecting the circular relationship among directions (**Fig. 4A, B**), consistent with prior reports for head-direction and stimulus orientation coding^43–45^.

**Fig. 4.**
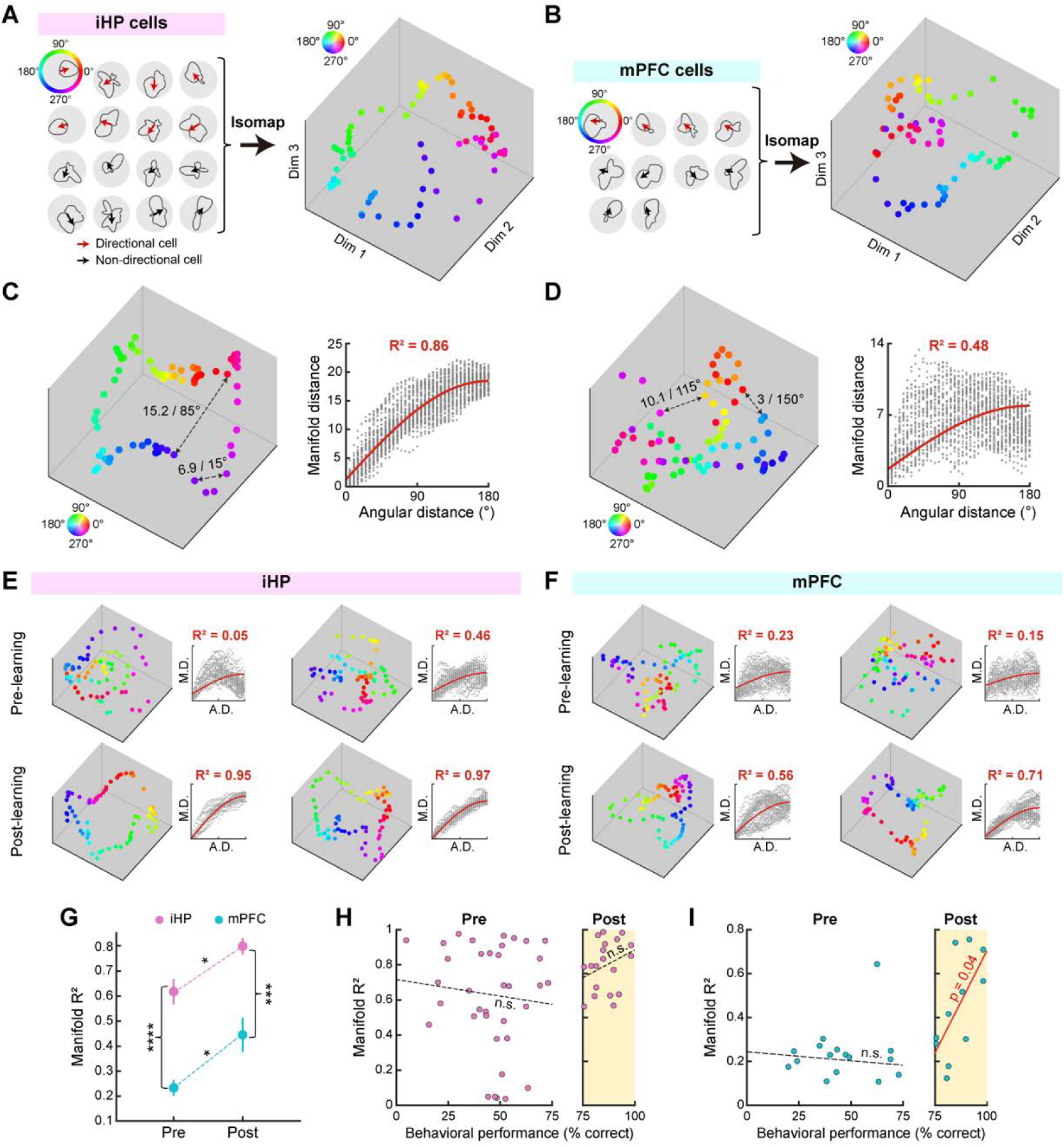
Directional neural manifolds in the iHP and the mPFC and their geometric structure over the course of learning. (**A**, **B**) Construction of directional neural manifolds from directional tuning curves of all simultaneously recorded neurons in the iHP (**A**) or the mPFC (**B**) using Isomap. Rose plots show spike distributions across VR direction, with arrows indicating the mean vector. (**C**) An example neural manifold with a robust ring-like structure (left). Numbers indicate the Euclidean distances in manifold space (i.e., manifold distances) and the angular distances between two data points connected by dotted arrows. The scatter plot (right) shows all pairwise manifold distances versus angular distances from the same manifold. The red sinusoidal regression fit was used to compute the R^2^ value, reflecting how strongly the manifold exhibits a ring-like structure. (**D**) Same as in **C,** but showing a weaker ring-like structure. (**E**, **F**) Additional examples of neural manifolds from the iHP (**E**) and the mPFC (**F**), and their sinusoidal fits relating pairwise manifold distances (M.D.) to angular distances (A.D.). The upper row shows pre-learning examples and the lower row displays post-learning examples. (**G**) Comparison of manifold R^2^ values between the iHP and the mPFC during pre- and post-acquisition stages. Data represent the mean ± SEM. (**H**) Scatter plot of behavioral performance versus manifold R^2^ for individual sessions, shown separately for pre- and post-learning stages. Dotted lines indicate non-significant linear regression fits. (**I**) Same as in **H**, but for the mPFC. The red line indicates a significant linear regression fit. *p < 0.05, ***p < 0.001, ****p < 0.0001; n.s., not significant.

To quantify the ring-like structure (i.e., circularity) of our manifolds, we assumed that each data point in a manifold should preserve its angular relationship in the three-dimensional embedded space. For example, in a ring-like manifold, two points representing neural states from distant directions should be separated by a large Euclidean distance in the manifold space, and nearby directions should be closer to each other (**Fig. 4C**). When this relationship breaks down, the manifold departs from a ring shape (**Fig. 4D**). We quantified this ring-like quality by fitting a sinusoidal regression to the pairwise Euclidean distances as a function of angular distances in the manifold space, using the variance explained (R^2^) as a metric for circularity.

We applied this method to evaluate the ring-like structures of the iHP (**Fig. 4E**) and mPFC (**Fig. 4F**) manifolds, comparing pre- and post-learning stages based on the manifold R^2^ values. The iHP manifolds exhibited significantly higher R^2^ values than the mPFC manifolds (F_(1,78)_ = 44.36, p < 0.0001; two-way ANOVA), indicating more robust ring-like structures (**Fig. 4G**). Additionally, supporting the results from the single-neuron analysis, R^2^ values significantly increased in the post-learning stage in both regions (F_(1,78)_ = 12.16, p < 0.0001, two-way ANOVA; iHP-Pre vs. iHP-Post, t_(53)_ = 2.43, p = 0.037; mPFC-Pre vs. mPFC-Post, t_(53)_ = 3.47, p = 0.0038; t-test with Bonferroni correction [n = 2]) (**Fig. 4G**). To determine whether this increase in R^2^ in post-learning reflected a gradual improvement in performance (i.e., an increase in the number of correct trials) or a discrete learning transition between pre- and post-learning stages, we analyzed the relationship between manifold R^2^ and trial correctness for each learning stage. In the iHP, R^2^ values showed no significant correlation with correctness in either stage (pre-learning, p = 0.59; post-learning, p = 0.26; robust linear regression) (**Fig. 4H**). In the mPFC, R^2^ values were not significantly correlated with correctness in pre-learning sessions (p = 0.6) but showed a significant correlation post-learning (p = 0.038; robust linear regression). These results suggest that the emergence of a robust ring-like structure in the manifolds reflects task learning rather than a mere increase in correct trials.

### Convergence of the iHP and mPFC manifolds to a shared geometry after learning

We investigated whether the enhanced circularity of post-learning manifolds resulted in greater similarity between the iHP and mPFC representations. We also checked whether these manifolds encoded reward location (West vs. East), consistent with the goal-biased tuning observed at the single-neuron level (**Fig. 3E–G**). We performed a decoding analysis of session type (West vs. East) based on the manifold structure. We quantified a manifold’s structure from the pairwise Euclidean distances between all of its data points by computing, for each session and region, a 72 × 72 matrix of Euclidean distances among its data points and vectorizing the upper triangle (n = 2556) to obtain a feature vector. These features were used for the subsequent decoding analysis.

We first asked whether West and East sessions formed separable clusters in this feature space. Multidimensional scaling (MDS) revealed that West and East sessions were not distinctly separated during pre-learning (**Fig. 5A**, left) but became more distinct during post-learning, with the iHP and mPFC manifolds clustering together within each condition (**Fig. 5A**, right), indicating similar geometry across regions. These observations justify the use of the pairwise distance features to decode the session type with a linear classifier.

**Fig. 5.**
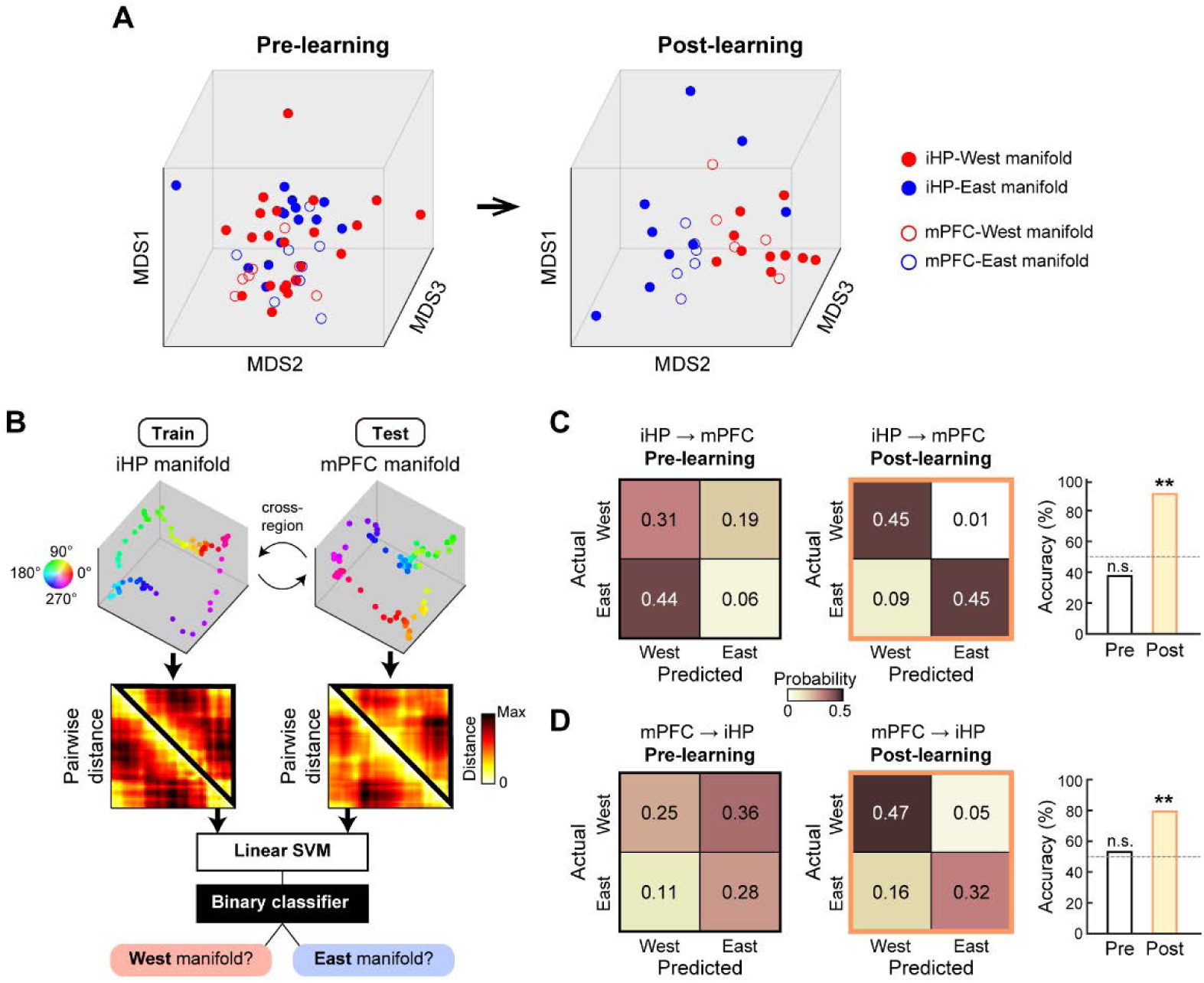
Geometric alignment of iHP and mPFC manifolds after learning. (**A**) Low-dimensional projections of individual iHP and mPFC neural manifolds from the West and East conditions using multi-dimensional scaling (MDS). Projections are shown separately for pre- and post-learning stages. (**B**) Schematic illustrating cross-regional decoding of the West and East conditions based on the structure of neural manifolds, estimated from pairwise Euclidean distances. iHP and mPFC manifolds are used interchangeably as training and testing sets. (**C**) Decoding results in pre- and post-learning stages for the iHP train–mPFC test condition, shown as a confusion matrix (left) and a decoding accuracy bar plot (right). The dotted line indicates chance-level decoding accuracy (50%). (**D**) Same as in **C**, but for the mPFC train–iHP test condition. **p < 0.01; n.s., not significant.

We employed cross-condition generalization performance (CCGP) analysis^22,46,47^ to decode session type (West vs. East) using a linear support vector machine (SVM) with the same feature vector. For example, we trained the SVM with feature vectors from iHP manifolds and tested it with feature vectors from mPFC manifolds to test whether features used to decode in one brain region could be generalized to the other (**Fig. 5B**). We analyzed pre- and post-learning sessions separately and decoding accuracy was quantified as the proportion of correctly labeled sessions. Decoding accuracy did not significantly differ from chance in pre-learning sessions (accuracy = 38% [6/16], p = 0.78) but was significantly above chance in post-learning (accuracy = 91% [10/11], p = 0.002; permutation test) (**Fig. 5C**). Training the SVM with mPFC manifolds and testing with the iHP manifolds yielded the same pattern, with accuracy not significantly exceeding chance in pre-learning sessions (accuracy = 53% [19/36], p = 0.28) but significantly above chance in post-learning sessions (accuracy = 79% [15/19], p = 0.005; permutation test). These findings indicate that after learning, manifold geometry encodes goal information as demonstrated by the successful decoding of West and East sessions. Furthermore, significant cross-region decoding shows that the geometric representations of the iHP and mPFC become aligned after learning.

### Differential temporal dynamics of manifold evolution between the iHP and the mPFC during learning

To examine the dynamics of manifold evolution on a trial-by-trial basis within a session, we defined “acquisition trial” as the one where the lower bound of the 90% confidence interval of the estimated learning curve crossed the chance level (50%). Based on the definition, we identified “acquisition sessions” as those in which (i) the acquisition trial occurred after the first 30 trials, and (ii) the mean accuracy after the acquisition trial exceeded 75% (**Fig. 6A**). We identified seven acquisition sessions, of which five had previously been labeled as pre-learning and the remaining two were post-learning.

**Fig. 6.**
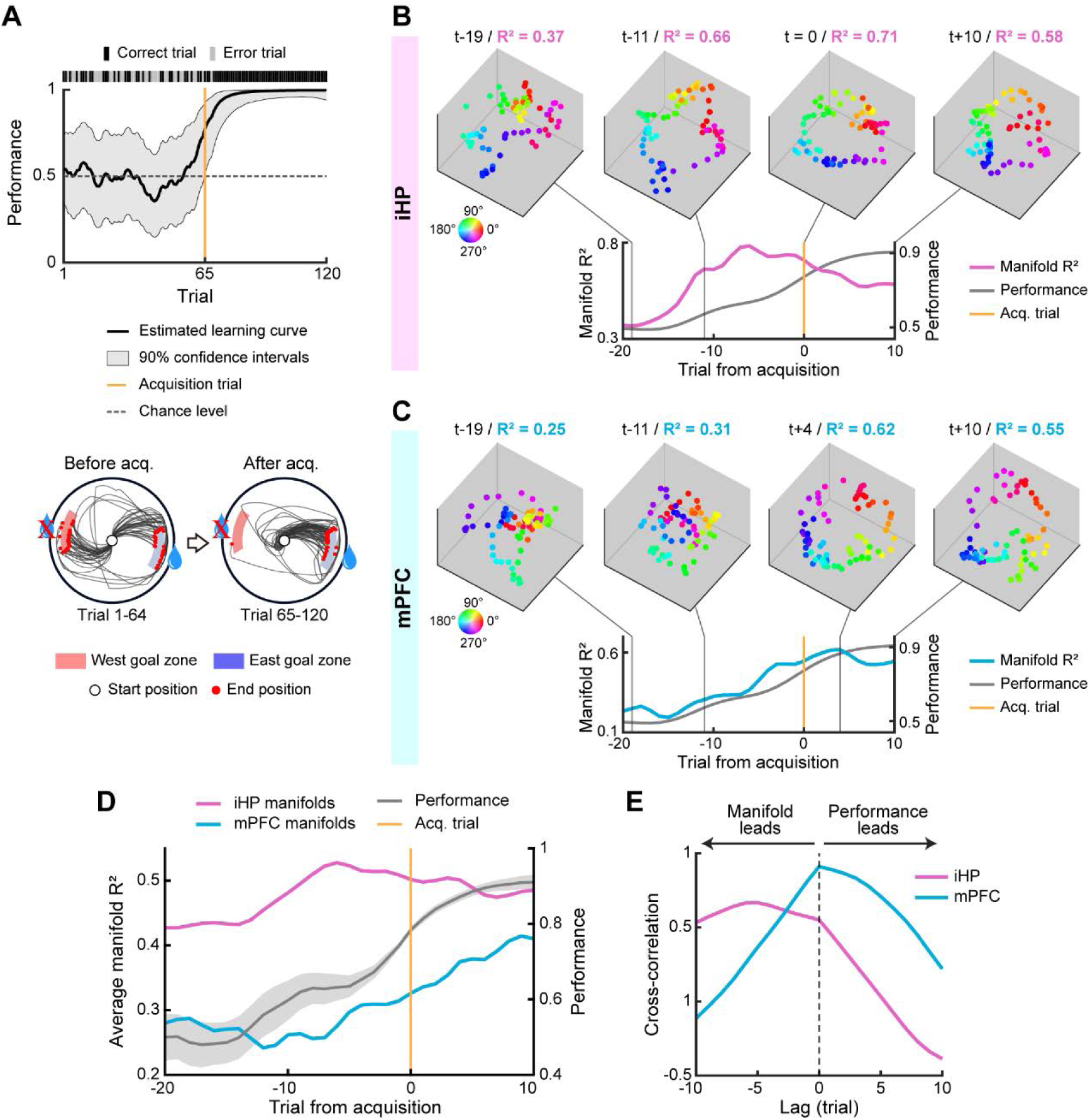
Across-trial dynamics of directional neural manifolds during learning. (**A**) Example behavioral data from an acquisition session showing within-session performance improvement, illustrated by an estimated learning curve (top) and trajectories before and after the acquisition trial (bottom). (**B**) Example of across-trial neural manifold dynamics in the iHP from one iteration of neuron subsampling across all acquisition sessions. Manifold examples are shown (top) with trial numbers relative to the acquisition trial and the corresponding manifold R^2^ above each example. Across-trial changes in manifold R^2^ values are shown with the average performance across acquisition sessions (bottom). Trial numbers are aligned relative to the acquisition trial, marked by a yellow vertical line. (**C**) Same as in **B**, but for the mPFC. (**D**) Average changes in manifold R^2^ around the acquisition trial in the iHP and the mPFC, shown together with performance. The average manifold R^2^ values were obtained by averaging across all iterations (n = 1000). (**E**) Cross-correlation between changes in manifold R^2^ and performance from **D**, separately for the iHP and the mPFC. Negative lags indicate that changes in manifold R^2^ precede changes in performance, and positive lags indicate the reverse.

Within these sessions, directional tuning curves were computed in a 20-trial sliding window in one-trial increments for all neurons recorded in the acquisition sessions. We then created directional manifolds for trials from −20 to +10 around the acquisition trial by randomly subsampling 20 neurons from each region within the acquisition sessions, aligned to the acquisition trial (t = 0). We calculated the manifold R^2^ across the aligned trials to evaluate the manifolds at the trial level. Representative examples from these subsamples exhibited distinct temporal dynamics between the two regions. That is, in an iHP example, the manifold started to exhibit a ring-like structure 11 trials prior to the acquisition trial (e.g., t-11) (**Fig. 6B**). However, in an mPFC example, the manifold did not exhibit a ring-like structure until the acquisition trial, and the manifold R^2^ peaked after the acquisition trial (**Fig. 6C**).

We examined the manifold R^2^ patterns around the acquisition trial by averaging the R^2^ values across all subsamples (**Fig. 6D**). Consistent with our previous results (**Fig. 4G**), average R^2^ values remained higher in the iHP than in the mPFC. Similar to the examples above (**Fig. 6B**, **6C**), in the iHP, manifold R^2^ increased abruptly before the acquisition trial and peaked 5 trials prior to the acquisition trial (**Fig. 6D**). In the mPFC, R^2^ values increased as performance improved around the acquisition trial, peaking after the acquisition trial. Cross-correlation analysis of manifold R^2^ and performance revealed that R^2^ changes in the iHP preceded performance (negative lag), while mPFC R^2^ changes occurred simultaneously with performance (zero lag) (**Fig. 6E**). These results suggest that the iHP develops a robust ring-like structure earlier than the mPFC, indicating a temporal precedence in learning-related changes in goal-directed navigation.

### Development of the mPFC manifolds via interactions with the iHP theta oscillations

We examined whether hippocampal-prefrontal interactions, specifically via theta oscillations (6-12Hz)^19,25,26,28,48,49^, contribute to manifold geometry changes. Local field potentials (LFPs) recorded alongside spikes revealed coherent theta oscillations in both regions (**Fig. 7A**). Many neurons in the iHP and mPFC were phase-locked to theta in the iHP or mPFC (**Fig. 7B**) or to theta oscillations from both regions (Rayleigh test, α = 0.01) (**Fig. 7C**), with the majority of neurons phase-locked to iHP theta (iHP neurons: pre-learning, 93.1%; post-learning, 94%; mPFC neurons: pre-learning, 61.5%; post-learning, 70.1%). For neurons phase-locked to both iHP and mPFC theta, we compared their preferred phases relative to iHP and mPFC theta; we observed that, for neurons in both the iHP and the mPFC, the preferred phase relative to iHP theta preceded that relative to mPFC theta (**Fig. 7D**). This temporal relationship was also evident in the examples in figures 7A and 7B. These temporal dynamics were observed regardless of the learning stage (**Fig. 7E**). These results suggest information flow from the iHP to the mPFC via the theta rhythm^25,26,50^.

**Fig. 7.**
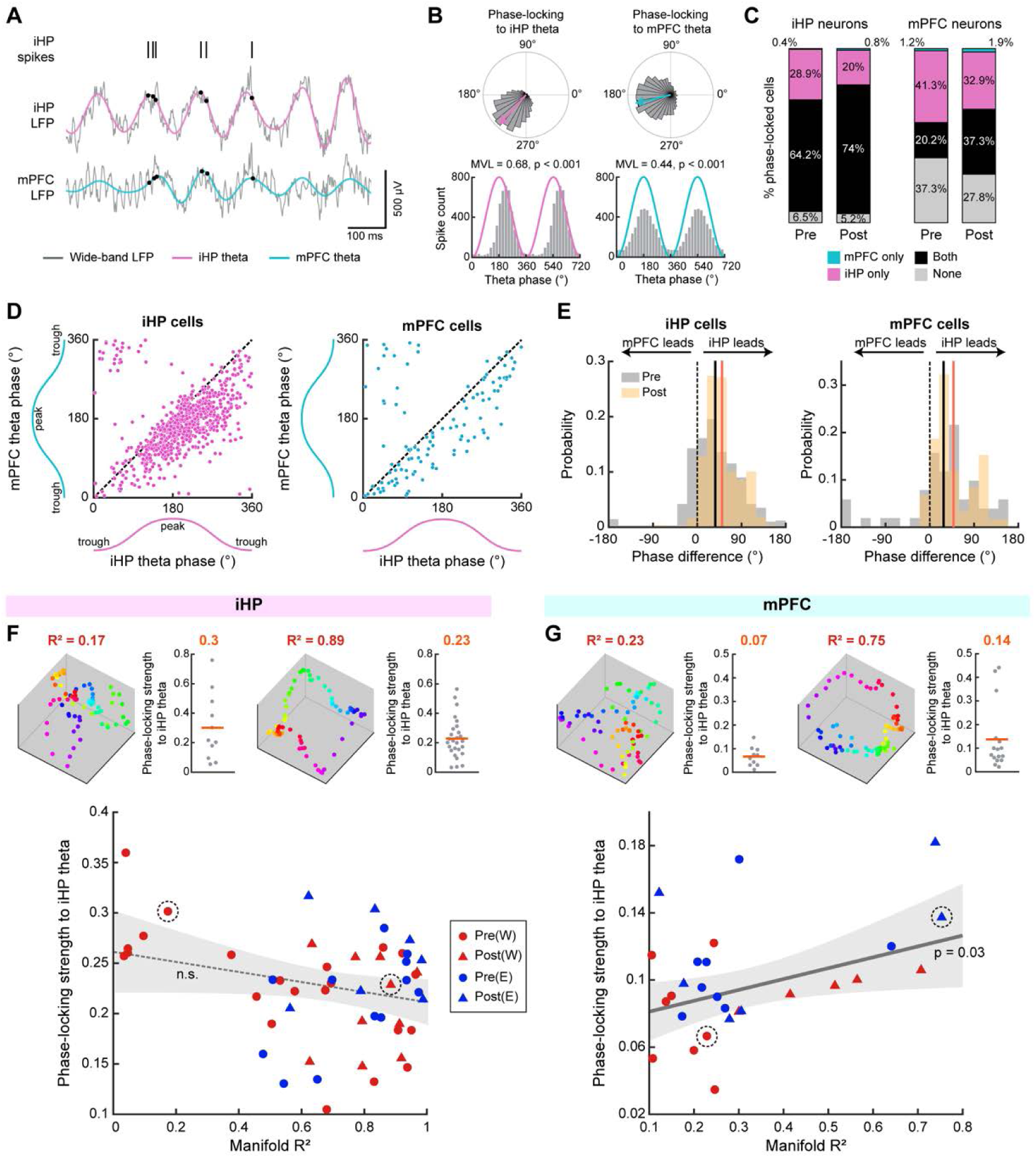
Theta phase-locking and its relationship to manifold geometry. (**A**) Representative spike trains from an iHP neuron, along with simultaneously recorded theta oscillations from the iHP and the mPFC. (**B**) Phase distributions of the neuron shown in **A** relative to iHP and mPFC theta oscillations. MVL, mean vector length. (**C**) Proportions of iHP and mPFC neurons phase-locked to iHP and mPFC theta oscillations. (**D**) Scatter plots of preferred phases relative to iHP and mPFC theta oscillations for iHP neurons (left) and mPFC neurons (right). (**E**) Histograms of phase differences between iHP and mPFC theta oscillations, shown separately for iHP (left) and mPFC neurons (right). Positive values indicate that iHP theta precedes mPFC theta. Black and orange vertical lines indicate average phase differences during pre- and post-learning stages, respectively. (**F**) Relationship between the ring-like structure of iHP manifolds and phase-locking strength to iHP theta oscillations. Example manifolds (highlighted with dotted circles in the scatter plot below) are shown above, along with their manifold R^2^ value (red) and average phase-locking strength to iHP theta (orange). Each gray dot represents a single neuron. The scatter plot below shows manifold R^2^ values and average phase-locking strength for all iHP manifolds. The dotted line indicates a non-significant linear regression fit, with the 95% confidence interval shaded in gray. (**G**) Same as in **F**, but for mPFC manifolds. The solid gray line indicates a significant linear regression fit.

The abovementioned results led us to hypothesize that the mPFC’s manifold geometry is influenced by theta oscillations from the iHP. We tested this hypothesis by examining the relationship between manifold R^2^ and the strength of phase-locking to iHP theta for the neurons comprising the manifolds (**Fig. 7F**). To compute the strength of theta phase-locking at the manifold level, we averaged the mean vector lengths from spike-phase histograms across all neurons used to construct the manifold (**Fig. 7C**). In the iHP, phase-locking strength to iHP theta showed no significant correlation with manifold R^2^ (Pearson’s r = −0.29; p = 0.072; robust linear regression) (**Fig. 7F**). In contrast, mPFC manifolds exhibited a significant positive correlation between phase-locking strength and manifold R^2^ (Pearson’s r = 0.42; p = 0.033; robust linear regression) (**Fig. 7G**). These findings suggest that theta-mediated interactions from iHP to mPFC play a critical role in shaping the geometric structure of mPFC neural manifolds during learning.

## Discussion

The current study investigated how task-related spatial representations dynamically changed in the iHP and the mPFC over the course of learning. Since it required several days to reach the learning criterion in our VR task, we were able to define pre- and post-learning sessions and directly compare neural activity between them. We observed significant directional tuning in both regions, with an increase in directional specificity after learning. We further confirmed this directional refinement by examining population-level representations with neural manifold analysis. Notably, post-learning manifolds from the iHP and the mPFC exhibited similar goal-informative geometry. When we examined the manifold dynamics at the trial level, iHP manifolds evolved earlier than mPFC manifolds. Finally, we discovered a consistent information flow from the iHP to the mPFC through theta oscillations; there was also a positive relationship between theta phase-locking and manifold geometry in the mPFC. Together, these results suggest that the iHP and the mPFC reshape their spatial codes during learning toward a shared, goal-informative geometry. Importantly, these learning-related changes were sequentially developed from the iHP to the mPFC, suggesting that the mPFC follows the spatial guidance of the iHP in goal-directed navigation.

### The presence of directional coding in the iHP and the mPFC and its relationship with place coding

Most rodent studies have emphasized positional correlates such as place cells when describing hippocampal-prefrontal spatial coding. However, in our 2D VR setting, we discovered that about 30% of iHP and mPFC neurons were directionally tuned, and their firing patterns could not be solely explained by the covariance between position and direction. These results are not entirely novel, considering previous findings that place cells can be modulated by head direction in radial arm maze and circular arena^51,52^. Additionally, it is noteworthy that, in a visually enriched 2D VR environment, dorsal hippocampus neurons were reported to be modulated by direction, even after minimizing the influence of position factor^40,41^. The authors explained that rodents heavily rely on allocentric visual cues to recognize space since other sensory cues (e.g., vestibular or olfactory) are not available in 2D VR environment. This may make the hippocampus more responsive to facing direction, similar to spatial view cells reported in the primate hippocampus^53,54^. Our results extend this phenomenon from the dorsal hippocampus to a broader network, including the iHP and mPFC.

The directional tuning we employed as the primary spatial correlate in our study shared several features with previously reported position coding in the hippocampal-prefrontal network. First, directional specificity was higher in the iHP than in the mPFC, consistent with lower proportions of position-selective cells and broader fields in the mPFC^19,21,22,42,55^. Second, many directional cells were biased toward goal-facing directions, similar to the overrepresentation of place coding near goal locations^36,56,57^. However, the relationship to task performance differs from that in classic place coding. Specifically, our data showed that directional tuning sharpened after learning in both iHP and mPFC, in agreement with another 2D VR study that showed positive correlation between the task performance and the proportion of direction-tuned cells^40^. By contrast, post-learning enhancement of hippocampal place coding has not been established. In the mPFC, there is evidence for a learning-related increase in positional information with improved performance^55^. Overall, directional tuning in our data captures key properties of well-known position coding, while enabling a measurement of learning-related changes in 2D VR across the hippocampal-prefrontal network.

We further characterized the directional tuning with neural manifold analysis. Since direction is a circular variable, we were able to evaluate the population-level representations in a simple and intuitive way based on the ring-like geometry of directional tuning, compared to manifold analysis of allocentric 2D positions^58,22,59^. We were able to identify ring-like geometry in both the iHP and the mPFC, as expected from directional tuning at the single-neuron level.

Crucially, the “ring-like” structure here did not represent an ideal, perfectly circular attractor as described in low-level sensory cortices or in canonical head-direction circuits^43,44^. Instead, the geometry reflected a goal-directed map since it carried information about the current goal location, as shown by successful decoding of session type from manifold features. Consistent with this observation, recent work has shown that hippocampal population geometry shifts from an unstructured to a more orthogonalized state that dissociates different task conditions after learning^60^. Accordingly, we interpret the strengthening of the ring-like structure post-learning not as a simple increase in directional precision across all 360°, but as the emergence of a more precise goal-oriented map of space for successful navigation.

### Sequential development of task-relevant spatial coding from the iHP to the mPFC

Previous studies on the hippocampal-prefrontal network have long asked what information flows from the hippocampus to the mPFC, and have searched for shared neural substrates across the two regions^42^. However, most prior studies have highlighted differences in spatially modulated firing between the two regions. For example, during free foraging, mPFC ensembles differentiated repeated exposures to the same environment, whereas hippocampal ensembles preserved spatial firing across exposure^61^. When rats performed the same W-maze alternation tasks in two different contexts, hippocampal neurons remapped in a context-specific manner, whereas prefrontal population activity generalized across contexts to represent the task structure^22^. Likewise, to the best of our knowledge, evidence that the two regions develop shared representations, especially through learning, has rarely been reported. Our data, in this regard, demonstrated that iHP and mPFC develop ring-like manifolds after learning, and that both manifolds carry goal information in their geometry as shown by cross-region decoding analysis. Therefore, our results provide compelling evidence that the hippocampus and mPFC indeed develop shared spatial representations as previously hypothesized.

We further investigated the temporal dynamics of these learning-related changes by evaluating how manifolds changed at the trial level as the rats’ performance increased sharply. We found that the ring-like structure emerged earlier in iHP manifolds than in the mPFC manifolds. These results align with the complementary learning systems account of the hippocampal-prefrontal network, which posits that the hippocampus is a fast-learning system and the prefrontal cortex is a slow-learning system^9,62^. In this framework, the hippocampus, long hypothesized to have a critical role in initial storage of memory, supports rapid acquisition of task-relevant information, whereas neocortical networks (especially the prefrontal cortex) integrate this information from the hippocampus to form a more generalized, schema-like representation through the memory consolidation process^4,9,10,62^. Although prior work mostly emphasized online coupling of the hippocampus and prefrontal cortex for successful behavior in well-trained rats^21,25,26^, our findings support the notion that these regions actually interact over longer timescales during long-term memory formation to develop coherent population-level geometry.

It is worth noting that most hippocampal-prefrontal research has focused on the dorsal hippocampus, even though its connections to the mPFC rely on indirect pathways such as the nucleus reuniens, subiculum, retrosplenial cortex^31,63,64^. By contrast, studies of the ventral hippocampus-mPFC pathway, which is based in direct connections, have mainly emphasized non-spatial domains such as anxiety and novelty^33,35^. The iHP-mPFC circuit has rarely been examined in the context of spatial learning, but a lesion study suggested that the intermediate hippocampus, rather than the dorsal or ventral hippocampus, is necessary for rapid (i.e., single-trial) learning in the water maze. The study proposed that the iHP-mPFC circuit is critical for the rapid translation of spatial encoding into behavior^65^. Our results may support this framework because prior work centered on the dorsal hippocampus did not show a comparable extent of convergence in spatial representations within the hippocampal-prefrontal network. Further studies are needed to establish whether these post-learning synchronization between the iHP and mPFC is unique to the iHP relative to dorsal or ventral-most part of the hippocampus through simultaneous recordings from all hippocampal subregions during learning.

### Hippocampal theta oscillation as a possible mechanism for the learning of spatial coding in the mPFC

Previous studies have demonstrated the importance of theta oscillations in the functional coupling of the hippocampus and the prefrontal cortex for successful memory-guided performance^19,25,28^. For example, in a Y-maze, it has been shown that theta coherence between the hippocampus and mPFC was selectively enhanced at the choice point, particularly when the rats were in a high-performance state^28^. In terms of spike phase-locking, mPFC neurons in a continuous spatial alternation task were more phase-locked to hippocampal theta in correct trials during the choice phase than in error trials or during forced choices. We also observed strong phase-locking of neurons in both the iHP and the mPFC to theta oscillations, although there was no clear enhancement of phase-locking after learning. Notably, we observed even stronger phase-locking to hippocampal theta in both regions (over 90% for iHP neurons and 70% for mPFC neurons) compared to the abovementioned studies focused on the dorsal hippocampus (< 80% for dorsal hippocampus units and < 60% in the mPFC). These results are consistent with previous reports showing stronger theta coherence between the intermediate/ventral hippocampus and the mPFC compared to the dorsal hippocampus^48^

Theta oscillations are thought to originate in the hippocampus and thus serve as a medium for transmitting information from the hippocampus to other cortical regions^12,66^. Indeed, hippocampal theta typically leads the mPFC in phase, as shown by the backward shifting of hippocampal theta oscillations to maximize covariance with mPFC theta^25,26,67^. Similarly, when we examined the preferred phases of spikes to iHP and mPFC theta oscillations, we found that the iHP theta preceded the mPFC theta in both pre- and post-learning stages. This allowed us to reconfirm the previously established hippocampus-to-prefrontal information transmission via theta oscillations. This also suggests that theta oscillations could be a crucial mechanism enabling the mPFC to acquire task-relevant spatial representations from the iHP.

However, it is noteworthy that studies on the relationship between theta oscillations and spatial representations in the hippocampus and prefrontal cortex have been scarce. It has been reported that incorporating hippocampal theta oscillations into the mPFC spiking activities increased the decoding accuracy of spatial position^21^. In another study, it has been shown that spatial decoding based on mPFC firing lags behind the hippocampus, and these temporal relationships are reinstated in theta-based coupling between the two regions^68^. However, to the best of our knowledge, there has been no empirical findings that directly investigated the dynamics of spatial representations in the mPFC along with theta-based coupling to the hippocampus. We demonstrated a positive relationship between the ring-like structure of mPFC manifolds and phase-locking strength to iHP theta of neurons comprising the manifold. These results may arise through enhanced coactivation of iHP-mPFC cell assemblies during phase-locked spiking to induce functionally similar spatial representations between the two regions^69^. Future work should aim to establish the causality of the theta coordination in reshaping spatial code in the mPFC by manipulating theta coupling and examining the following geometric changes during spatial learning^70^.

## Methods

### Subjects

Six male Long-Evans rats (10 weeks old) were obtained and used in this study. Animals were individually housed in a temperature- and humidity-controlled room under a 12-hour light/dark cycle (lights on at 8:00 AM). All behavioral training and electrophysiological recordings were conducted during the light phase. Rats had ad libitum access to food, but water was restricted to maintain body weights at 85% of free-feeding weight at the start of the experiment. All experimental procedures were approved by the Institutional Animal Care and Use Committee of Seoul National University (SNU-200504-3-1).

### 2D VR apparatus

We developed a rodent VR apparatus designed for 2D navigation, as previously described^39^.The setup consisted of multiple LCD monitors, a spherical treadmill, rotary encoders, a body-restraint system, and a reward delivery system (**Fig. 1A**). Five adjacent LCD monitors, covering 270° of the visual field, were used to present the VR environment. To provide a realistic visual experience, we used a high-resolution game engine (Unreal Engine 4.14.3; Epic Games) to create the VR environment, which consisted of a circular arena surrounded by multiple distal landmarks (**Fig. 1C**). Rats were body-restrained with a custom-made body jacket that was fixed to an aluminum profile structure, which secured the animal’s body above the spherical treadmill. The spherical treadmill consisted of a silicone-coated Styrofoam ball (400 mm in diameter) supported by multiple ball bearings. As the rats rolled the ball, their movement was tracked by three rotary encoders (DBS60EBGFJD1024; Sick) attached to the surface of the ball. The encoder signals were then transmitted to the computer via an Arduino Leonardo, and synchronized with the animal’s movement in the VR environment using MATLAB R2021a (MathWorks) and Unreal Engine. The reward delivery system included a licking port placed in front of the animal, which rotated in accordance with the animal’s orientation. The port remained retracted during navigation but was extended toward the snout by a linear actuator (L16-R; Actuonix Motion Devices) upon the rat’s successful entry into a reward location. An infrared sensor (FD-S32; Panasonic) detected the rats’ licking behavior to trigger the solenoid valve (VA212-3N; Aonetech) to dispense a water reward (10 µl). Reward delivery was also controlled by Unreal Engine via the Arduino Uno interface.

### Behavioral paradigm

After several days of handling, rats were introduced to the VR apparatus (“Habituation”) and then learned to navigate the VR environment by rolling the spherical treadmill. During this phase (“Beacon-based navigation task”), rats received a water reward upon reaching a flickering checkerboard-patterned sphere that appeared at random locations within a circular arena (120 cm in diameter). Animals that completed more than 40 trials in 60 minutes on two consecutive days were considered proficient in navigating the VR space and advanced to pre-surgical training of the main task.

In the main task (“Goal-directed spatial navigation task”), rats navigated the same circular arena to find a hidden reward location based on allocentric visual cues (**Fig. 1B-D**). Two unmarked goal zones were located along the perimeter, positioned symmetrically on the west (170°) and east (350°) sides. One of these zones served as the reward location, while the other was non-rewarding. Both zones were placed slightly inward from the arena boundary to prevent wall-following behavior. Each trial began with the rat positioned at the center of the arena, facing either north (90°) or south (270°), chosen pseudorandomly. Trials ended when the rat entered either goal zone or after 90 s had elapsed, but only entry into the designated reward location resulted in reward delivery.

Training began with the West condition, in which the West goal zone was rewarded. Once rats achieved >75% correctness on two consecutive days, the reward location was reversed to the East and the West zone became non-rewarding (i.e., the East condition). Rats acquired the new reward location through trial and error. After completing pre-surgical training in both West and East conditions, rats underwent surgical implantation of a hyperdrive (see below). Following a one-week recovery period, animals resumed task training with the same sequence: beacon-based navigation followed by goal-directed spatial navigation in the West and East conditions.

All behavioral and neural data analyses were conducted within these post-surgical West and East sessions. To define learning stages based on performance, sessions with <75% accuracy were classified as pre-learning, and those with >75% accuracy were defined as post-learning. See figure 1D for the full experimental timeline.

### Hyperdrive implantation

To record single-unit activity and local field potentials (LFPs), a hyperdrive containing 24 independently movable tetrodes was built in-house. Each tetrode was assembled from four strands of formvar-insulated nichrome wire (17.8-µm diameter), thermally bonded and gold-plated using a Nano-Z system (Neuralynx) to reduce impedance to ∼200 kΩ at 1 kHz. The hyperdrive had two separate bundles: one targeting the medial prefrontal cortex (mPFC; 8 tetrodes), and the other targeting the intermediate hippocampus (iHP; 16 tetrodes).

To surgically implant the hyperdrive, rats were anesthetized with an intraperitoneal injection of sodium pentobarbital (Nembutal, 65 mg/kg), and placed in a stereotaxic frame (Kopf Instruments). Anesthesia was maintained throughout the procedure with 0.5-2% isoflurane. Following scalp incision, two craniotomies were made at coordinates targeting the mPFC (3.3 mm anterior to bregma, 1.3 mm lateral to the midline) and iHP (5.9 mm posterior to bregma, 5.9 mm lateral to the midline), each sized to accommodate the corresponding bundle. The hyperdrive was then slowly lowered to the brain surface and secured to the skull using 11 anchor screws and bone cement. A ground wire was connected to a skull screw placed above the cerebellum. Postsurgical care included oral administration of ibuprofen syrup and overnight monitoring in an intensive care unit.

### Electrophysiological recording

After allowing the rats seven days to recover from surgery, individual tetrodes were gradually lowered to target the mPFC and the iHP. We aimed to lower the tetrodes to their target depths by the first day of the goal-directed spatial navigation task. Neural signals were recorded using a Digital Lynx acquisition system (Neuralynx). Signals were amplified 1000–10,000-fold and bandpass filtered between 300 and 6000 Hz for spike detection. Spike waveforms exceeding a manually adjusted threshold (typically 60–100 µV) were digitized at 32 kHz and timestamped.

Behavioral data–including trial events, allocentric position, and facing direction, and other task-related variables–were synchronized with neural recordings via TTL signals generated by the VR system at 30 Hz. All data were collected during the post-surgical West and East sessions.

### Histology

After all recording sessions were completed, the tip of each tetrode was marked with electrolytic lesions using a weak current (10 μA for 10 s)., rats were sacrificed with an overdose of CO_2_ and transcardially perfused, first with phosphate-buffered saline and then with a 4% (v/v) formaldehyde solution. The brain was extracted and stored in a 4% (v/v) formaldehyde–30% sucrose solution at 4°C until it sank to the bottom of the container. It was subsequently coated with gelatin, soaked again in 4% (v/v) formaldehyde–30% sucrose solution. The brain was sectioned in the coronal plane at 40-μm thickness using a freezing microtome (HM 430; ThermoFisher Scientific). All sections containing the mPFC and iHP were collected and stained with thionin. Photomicrographs of each brain section were acquired using a microscope equipped with a digital camera (Eclipse 80i; Nikon). To estimate the positions of the tetrodes, the configuration of each bundle was reconstructed based on histology results, and compared with the original configuration to determine tetrode identity (Voxwin, UK).

### Unit isolation

All single units were manually isolated using a custom program (WinClust), as previously described. Various waveform parameters (i.e., peak amplitude, energy, and peak-to-trough latency) were used to isolate single units, but peak amplitude was the primary criterion. Units were excluded if more than 1% of spikes occurred within the refractory period (1 ms) and the mean firing rates during the task epoch (from trial start to goal arrival) were lower than 0.5 Hz.

### Single-unit analysis

#### Cell filtering

Single units recorded from both pre- and post-learning sessions in the West and East conditions were included in the analysis. Only units from sessions with more than 50 valid trials, after excluding void trials caused by the experimenter’s intervention or mechanical issues, were considered. Units were further classified into putative excitatory and inhibitory neurons based on their mean firing rate and spike width: neurons with a mean firing rate > 10 Hz and spike width < 300 μs were classified as inhibitory. All subsequent analyses were conducted within putative excitatory neurons recorded from the iHP (n = 1228) and mPFC (n = 410). We only analyzed spikes during active movement (speed > 5cm/s, as measured by rotary encoders).

#### Directional tuning

Directional tuning was assessed based on the allocentric facing direction of the virtual agent in the VR environment (referred to as VR direction). For each putative excitatory neuron, a directional tuning curve (spike-direction histogram) was constructed by dividing the number of spikes in each 1° direction bin by the time the animal spent in that bin. We calculated the mean vector length from the directional tuning curve as a metric for the specificity of directional tuning. To define directional cells, we computed a surrogate distribution of mean vector length by shuffling spike times (n = 1000). We then compared this distribution to the actual mean vector length (α < 0.05). For visualization and subsequent analyses, directional tuning curves were Gaussian-smoothed (σ = 30°).

#### Spatial tuning

To create a spatial rate map, the circular arena was divided into 24 × 24 bins (bin size = 5 cm) and the number of spikes was divided by the duration of occupancy to obtain the firing rate of each spatial bin. The map was then smoothed by Skaggs’ adaptive binning method^71^. We computed the spatial information score with the smoothed rate map using the following equation:

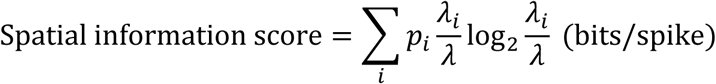

where *i* denotes bin, *p_i_* is the occupancy probability in the *i*th bin, *λ_i_* is the mean firing rate in the *i*th bin, and *λ* is the overall mean firing rate.

#### Goal-tuned cells

For each directional cell, the preferred direction was defined as the peak firing direction in the smoothed directional tuning curve. A neuron was classified as goal-tuned if its preferred direction was within a 90° window centered on the rewarding goal zone. In the West condition, the goal-centered window was defined as 170° ± 45°, and in the East condition as 350° ± 45°.

### Neural manifold analysis

#### Constructing directional neural manifolds

To capture population-level representations of directional tuning, we constructed directional neural manifolds separately for the iHP and the mPFC in each recording session. Only sessions with at least eight simultaneously recorded neurons from the same region were used to construct neural manifolds (iHP, n = 55; mPFC, n = 27). For each neuron, a directional tuning curve was computed using 5° bins (72 bins covering 360°), Gaussian-smoothed (σ = 30°), and was z-scored. Tuning curves of all simultaneously recorded neurons were concatenated to generate a 72 × *n* population activity matrix, where *n* is the number of neurons. We then applied isometric feature mapping (Isomap; k = 15) to this high-dimensional activity matrix to reduce the dimensionality to three, resulting in a 3D manifold consisting of 72 points.

#### Evaluating the manifold geometry

To quantify the ring-like geometry of the directional manifold, we compared the Euclidean distances between all pairs of 72 points in the 3D manifold (i.e., manifold distance), to the corresponding angular distances as defined in the VR space. A ring-shaped manifold is expected to show a positive relationship between manifold distance and angular distance. We therefore applied a sinusoidal regression of pairwise manifold distances as a function of angular distances, as follows:

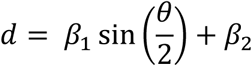

where *d* is the Euclidean distance between points, *θ* is the angular distance (in radians) between them, and the equation models the relationship for a given manifold. The R^2^ values from this regression (referred to as manifold R^2^) were used as the primary metric to quantify how closely the manifold approximates a ring-like geometry. Higher R^2^ values indicate that the low-dimensional projections preserve the circular organization of directional coding.

#### Manifold-based decoding of session type

To test whether the geometry of directional neural manifolds carried information about the current reward location (West vs. East), we performed a decoding analysis using pairwise Euclidean distances of the manifolds as features. For each manifold, we computed a 72 × 72 pairwise distance matrix and vectorized the upper triangle (2556 features) to obtain a feature vector. We first applied multidimensional scaling (MDS) to visualize potential clustering of manifolds by session type. The 2556 feature vectors from both regions were reduced and projected into three-dimensional space, separately for pre- and post-learning stages. We then performed session type decoding with linear support vector machine (SVM) using the same pairwise distances as feature vectors (*fitclinear* function in Matlab).

Decoding was performed in two ways: within-region and cross-region. Within-region decoding was performed separately for the iHP and mPFC. For each learning stage (pre- and post-), feature vectors from West and East conditions were combined and labeled. A linear SVM was trained and tested using leave-one-out cross-validation to classify West and East conditions. The decoding accuracy was computed as the fraction of correctly labeled sessions. Statistical significance was assessed with a 1,000-iteration permutation test, in which session labels were shuffled. Cross-region decoding was conducted to test whether both regions shared similar geometry when classifying West and East conditions. Here, the SVM was trained on features from one region (e.g., iHP) and tested on the other (e.g., mPFC) within the same learning stage, and vice versa. Accuracy and p-values were computed as in the within-region decoding.

#### Across-trial manifold dynamics

To capture across-trial dynamics of directional manifolds during learning, we focused on the acquisition sessions, which exhibited rapid within-session improvement in performance. For each session, we estimated a learning curve using a Bayesian state-space model, as previously described^36,72,73^. We defined an acquisition trial as the trial whose lower bound of 90% confidence interval of the learning curve crossed the chance level of 50%. A session was defined as an acquisition session if (i) the acquisition trial occurred after the first 30 trials and (ii) the mean accuracy after the acquisition trial exceeded 75%. Using all neurons recorded in the iHP and mPFC, we computed trial-by-trial directional tuning curves with a 20-trial window. To construct across-trial directional manifolds, we randomly subsampled 20 neurons (per iteration) from each region, aligned the trials relative to the acquisition trial (from −20 to +10), built directional manifolds using the trial-by-trial directional tuning curves as described above, and computed R^2^ values for each aligned trial. We performed 1,000 iterations and averaged the resulting R^2^ values across iterations to obtain the mean trajectory of R^2^ around the acquisition trial.

### Local field potential analysis

#### Preprocessing

Raw local field potentials (LFPs) were downsampled from 32 kHz to 2 kHz (Rate Reducer; Neuralynx). Theta oscillations (6–12 Hz) were extracted with a zero-phase, third-order Butterworth band-pass filter (*filtfilt*, MATLAB). To select a representative channel in each session, we computed the power spectral density using *mtspectrumc* in Chronux Toolbox and calculated the average theta power during behavioral task epochs. For the iHP, only tetrodes positioned in CA1 were considered (excluding CA2/CA3). For each region (the iHP and the mPFC), the channel with the highest average theta power was chosen as the representative channel. We then visually inspected the raw LFPs from that channel to assess for excessive noise. If the selected channel was noisy, the channel with the next highest power was chosen instead.

#### Theta phase-locking

Using theta from the representative channel, instantaneous theta phase was computed using the Hilbert transform. Theta phase of each spike was estimated by nearest-neighbor interpolation. Phases were expressed with 0° at the trough and 180° at the peak. We defined theta phase-locked neurons by testing non-uniformity of spike-phase distributions with the Rayleigh test (α < 0.01), computed separately for iHP and mPFC theta oscillations. The preferred phase of each unit was the circular mean (μ) from the Rayleigh test. For units recorded in sessions with simultaneous iHP and mPFC LFPs (iHP, n = 1130; mPFC, n = 410), we quantified the theta phase difference by subtracting the preferred phase relative to mPFC theta from that relative to iHP theta. To relate phase-locking to directional manifold geometry, we quantified each manifold’s phase-locking strength by averaging the mean vector length across units used to construct that manifold. We then examined the relationship between manifold R^2^ and phase-locking strength using robust linear regression.

## Acknowledgments

This work was supported by the National Research Foundation of Korea (RS-2025-02303740, RS-2024-00452391), and Mid-Career Bridging Program through Seoul National University for Inah Lee.

